# The relationship between fasting-induced torpor, sleep and wakefulness in the laboratory mouse

**DOI:** 10.1101/2020.05.05.076570

**Authors:** Yi G. Huang, Sarah J. Flaherty, Carina A. Pothecary, Russell G. Foster, Stuart N. Peirson, Vladyslav V. Vyazovskiy

**Affiliations:** Department of Physiology, Anatomy and Genetics, University of Oxford, Parks Road, Oxford, OX1 3PT, United Kingdom; Sleep and Circadian Neuroscience Institute, Nuffield Department of Clinical Neurosciences, Oxford Molecular Pathology Institute, Sir William Dunn School of Pathology, South Parks Road, Oxford OX1 3RE, United Kingdom

## Abstract

Torpor is a regulated reversible state of metabolic suppression used by many mammalian species to conserve energy. Although torpor has been studied extensively in terms of general physiology, metabolism and neuroendocrinology, the effects of hypometabolism and associated hypothermia on brain activity and states of vigilance have received little attention. Here we performed continuous monitoring of electroencephalogram (EEG), electromyogram (EMG) and peripheral body temperature in adult, male C57BL/6 mice over consecutive days of scheduled restricted feeding. All animals showed prominent bouts of hypothermia that became progressively deeper and longer as fasting progressed. EEG and EMG were markedly affected by hypothermia, although the typical electrophysiological signatures of NREM sleep, REM sleep and wakefulness allowed us to perform vigilance-state classification in all cases. Invariably, hypothermia bouts were initiated from a state indistinguishable from NREM sleep, with EEG power decreasing gradually in parallel with decreasing body temperature. Furthermore, during deep hypothermia REM sleep was largely abolished, but we observed brief and intense bursts of muscle activity, which resembled the regular motor discharges seen during early ontogeny associated with immature sleep patterns. We conclude that torpor and sleep are electrophysiologically on a continuum, and that, in order for torpor to occur, mice need to first transition through euthermic sleep.

## Introduction

States of vigilance in mammals are traditionally defined based on behavioural criteria and brain activity.^1–3^ Sleep is typically defined as a state of relative immobility and reduced sensory responsiveness, while wakefulness is characterised by movement and active engagement with the environment. These characteristics of an awake state are thought to be essential for its main functions, including feeding, mating or defence against predation. Whilst the relevance of immobility and sensory disconnection for sleep functions remains a matter of debate, it is likely that immobility is essential for the efficient execution of information processing or other essential functions related to energy and cellular homeostasis ^4,5^. The temporal pattern of vigilance states in most animals represents a balance between the competing needs to stay awake and sleep, which varies between species, between individuals of the same species and across ontogeny.^6,7^ Sleep is timed by an endogenous circadian clock and a homeostatic drive for sleep which builds during wake. These two processes allow the alignment of numerous aspects of behaviour and physiology with the occurrence of ecological factors such as light, food availability, ambient temperature and risk of predation.^8,9^

Sleep and wakefulness are distinguished by characteristic patterns of brain activity (an electroencephalogram or EEG), which arise from a dynamic interplay among numerous cortical and subcortical sleep-wake controlling circuits. During wakefulness, cortical activity is characterised by fast, a low amplitude EEG dominated by activities in a theta-frequency range (6–9 Hz), whilst non-rapid eye movement (NREM) sleep is dominated by slow waves (typically 0.5–4 Hz), arising within thalamocortical networks.^1,10^ The amplitude of slow-waves during NREM sleep is an established marker of the homeostatic sleep drive, which increases with prolonged wake and decreases with sleep.^11^ By contrast, another state of sleep - rapid eye-movement (REM) sleep – is typically characterised by lower amplitude, higher frequency, and theta-frequency oscillations, i.e. similar to the waking state.^1^

The sub-division of vigilance states into wake and sleep based on EEG and electromyography (EMG) signals and behaviour is relatively straightforward under physiological conditions, such as when the body temperature is normal (euthermic). However, significant deviation from normal body temperature may result in the occurrence of states that do not satisfy the criteria for wake and sleep established for euthermic conditions. One of the most notable states is torpor, defined as a regulated and reversible state of metabolic suppression.^1,12,13^ Torpor is used by many mammals as a strategy to conserve energy, particularly during periods of perceived or actual food shortage.^13^ Like sleeping mammals, torpid mammals appear quiescent, have reduced mobility and decreased responsiveness to sensory stimuli. However, unlike sleeping mammals, torpid mammals experience marked decreases in metabolic status, heart and breathing rates (sometimes down to just 1% of baseline values), and in body temperature (down to near ambient temperature).^14^ States of vigilance are typically defined based upon brain activity and behaviour. However, definitions of torpor lack such correlates and have been defined using systemic changes, primarily metabolic rate and/or body temperature. This raises the critical question of whether, and to what extent, sleep and torpor are similar or divergent states. The central aims of this paper are to address this question.

There have been relatively few studies in which EEG was recorded during torpor. In such cases torpor has been defined, rather arbitrarily, as a reversible decrease in T_core_ below 30 °C and a metabolic rate of 25% below basal metabolic rate (BMR).^15^ These studies were on seasonal torpor in the Arctic ground squirrel and daily torpor in the Djungarian hamster.^16,17^ The EEG during torpor in seasonally hibernating animals is enriched with slow-wave activity (SWA), not dissimilar to those during NREM sleep. However, these slow-waves are heterogeneous in their characteristics: at the start of torpor they cannot be differentiated from those that are typical of NREM sleep; as torpor progresses and T_core_ decreases further, the amplitude of these slow waves decreases markedly and their spectral peak shifts towards slower frequencies.^16,17^ These changes are likely to be at least partly due to a thermodynamic effect: lower temperatures slow the rates of all biochemical reactions including those occurring within neurons and at synapses.^18–21^ However, they could also be due to cooling-induced changes on synaptic connections, which have been reported in several species undergoing both natural and pharmacologically-induced torpor.^22–24^

Studies on Arctic ground squirrels and Djungarian hamsters has also yielded interesting insights into the interaction and possible functional differences between torpor and sleep. For example, immediately after Djungarian hamsters and Arctic ground squirrels emerge from a bout of torpor into euthermia, they go into a deep sleep characterised by significantly elevated SWA. This observation led to the hypothesis that during torpor animals are deprived of sleep.^16,25^ One interpretation of these findings is that, although torpor and sleep share similarities in terms of EEG activity, the former does not fully overlap with the latter functionally.^26,27^ Another possibility is that sleep, wake and torpor are not necessarily mutually exclusive states: animals can potentially exhibit not only sleep-like but also awake states during torpor.

In this study, we performed continuous monitoring of EEG, EMG and surface body temperature (T_surface_) in mice undergoing fasting-induced torpor. Although EEG has been studied during torpor in mammals that undergo hibernation (e.g. the Arctic ground squirrel) and those that undergo daily torpor (e.g. the Djungarian hamster),^16,25^ torpor has been largely ignored in the laboratory mouse, which readily enters daily torpor as a result of fasting.^28,29^ We observed that fasting-induced torpor is invariably entered through a state electrophysiologically indistinguishable from NREM sleep, but EEG power (the strength of a specific frequency in the signal) decreases drastically as a function of hypothermia. Moreover, deep hypothermia was accompanied with conspicuous intense bursts of EMG activity occurring regularly, especially at lowest body temperatures. We conclude that wakefulness, sleep, and torpor are likely not mutually exclusive states, but that sleep and torpor are continuous and share important similarities in terms of their electrophysiological characteristics. Furthermore, our findings have yielded the possibility that hypometabolism may unmask an underlying brain state that resembles a pattern of activity typical during early ontogeny.

## Materials and Methods

### Animals and recording conditions

Adult, male C57BL/6J mice were used in this study (Charles River; n = 6; aged 12 weeks). Throughout the experiment, mice were individually housed in custom-made clear plexiglass cages (20 x 30 x 35 cm) on a 12:12 h light-dark (12:12 LD) cycle for the duration of the experiment (**Fig. 1a**) inside sound-attenuated, ventilated recording chambers (Campden Instruments, Loughborough, UK; two cages per chamber). Each chamber was illuminated at approximately 200 lux by a warm white LED strip lamp during the light phase of the 12:12 LD cycle. Room temperature and relative humidity were regulated at 20 ± 1 °C and 60 ± 10%, respectively. T_surface_ was continually recorded using thermal imaging cameras (Optris Xi80, Berlin, Germany) (**Fig. 1a**). *Ad libitum* water was provided throughout the study. All procedures were performed in compliance with the United Kingdom Animals (Scientific Procedures) Act of 1986, as well as the University of Oxford Policy on the Use of Animals in Scientific Research (PPL P828B64BC). All experiments had approval from the University of Oxford Animal Welfare and Ethical Review Board.

**Figure 1.**
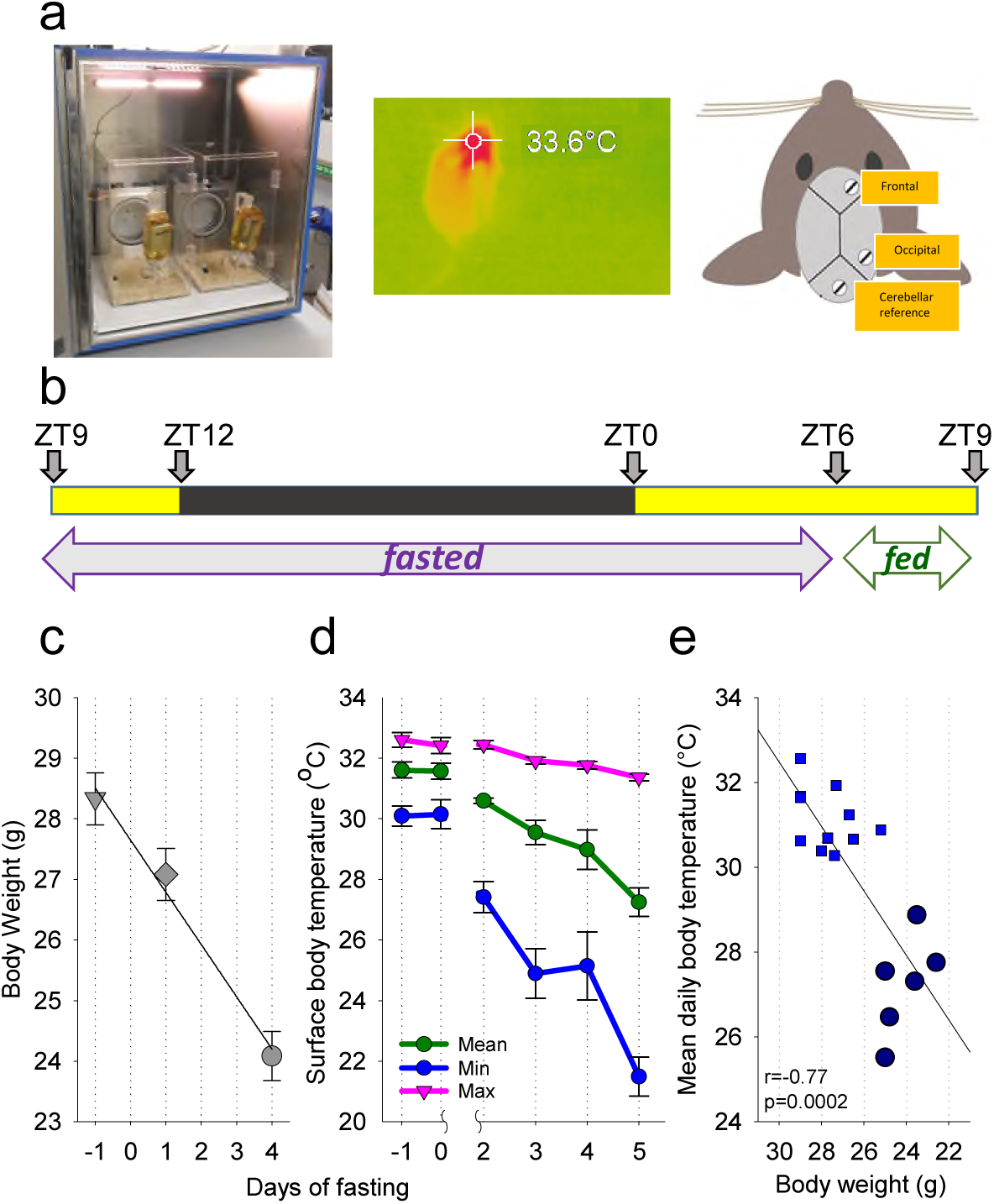
The experimental design and effects of food restriction on body weight and surface body temperature. **a**, Left: photograph showing the recording chamber with two Plexiglas cages for individually housed mice. Middle: A photograph of the thermal imaging camera and a representative thermal image of a mouse acquired with the camera. Right: schematic diagram showing the position of EEG electrodes. **b**, Experimental design illustrating the timing of feeding and fasting relative to the LD cycle. **c**, The time course of body weight across the experiment. Mean values (n=6, SEM). **d**, The time course of peripheral body temperature, shown separately for the maximum, mean and minimum daily temperature values, irrespective of the time of day or behavioural state. Mean values (n=6, SEM). **e**, The relationship between body weight and mean daily surface body temperature. The data points correspond to individual animals. Each animal contributes with three data points, corresponding to the days when body weight was measured (The data for Day 5 of fasting are depicted as large dark blue circles).

### Surgical procedure and experimental design

Animals underwent cranial surgery to implant custom-made EEG and EMG headmounts as described previously.^30–32^ Each headmount consisted of three stainless steel screw EEG electrodes (SelfTapping Bone Screws, length 4 mm, shaft diameter 0.85 mm; InterFocus Ltd, Cambridge, UK) and two stainless steel EMG wires, all attached to an 8-pin surface mount connector (8415-SM, Pinnacle Technology Inc, Kansas, USA). Surgical procedures were carried out using aseptic technique under isoflurane anaesthesia (5% for induction; 1.5–2.5% for maintenance). Animals were head-fixed during surgical procedures using a stereotaxic frame (David Kopf Instruments, California, USA).

Viscotears liquid gel (Alcon Laboratories Limited, Hemel Hempstead, UK) was applied at regular intervals to protect the eyes. Two headmount screws were implanted epidurally over the frontal (M1 motor area, anteroposterior (AP) +2 mm, mediolateral (ML) 2 mm) and occipital (V1 visual area, AP -3.5–4 mm, ML +2.5 mm) cortical areas (**Fig. 1a**). The third screw acted as a reference electrode and was implanted over the cerebellum; additionally, an anchor screw was implanted contralaterally to the frontal screw (with the tip within the cranium) to stabilise the head implant. Two stainless steel wires were inserted either side of the nuchal muscle for recording EMG. All headmount screws and wires were stabilised using dental cement (Associated Dental Products Ltd, Swindon, UK). Overall, this configuration gave two EEG derivations (frontal vs. cerebellum and occipital vs. cerebellum) and one EMG derivation. All animals were given subcutaneous (s.c.) normal saline and maintained on thermal support throughout surgery and for the subsequent 1–2 h. Analgesics were administered pre-and post-operatively (meloxicam 1–2 mg/kg, s.c., Metacam, Boehringer Ingelheim Ltd, Bracknell, UK). A 7-day recovery period was permitted prior to cabling the animals for recording. Mice were habituated to the recording cable for 2 days before recordings were used in analyses.^22-24^

### Restricted feeding paradigm

A restricted feeding (RF) paradigm, partly based upon a previous protocol, was used.^32^ Recordings began at ZT9 (Zeitgeber time; ZT0 = lights on, ZT12 = lights off). After obtaining two stable baseline 24h recordings with food provided *ad libitum* (defined from here as Days -1 and 0 of fasting), food was removed at ZT9 and subsequently made available to the animals only between ZT6-9 each day (**Fig. 1b**). This paradigm was chosen because, as demonstrated previously, it results in an occurrence of hypothermic bouts.^32^ Animals were weighed at ZT6 on Days -1, 2 and 5. The experiment was terminated at the end of Day 5 and food was subsequently provided *ad libitum*.

### Signal processing

EEG data was acquired using the Multi-channel Neurophysiology Recording System (TDT, Alachua FL, USA), as previously.^10,21,22,34^ EEG and EMG data were sampled at 256.9 Hz (filtered between 0.1–100 Hz), amplified (PZ5 NeuroDigitizer pre-amplifier, TDT Alachua FL, USA) and stored on a local PC. Data were resampled offline at 256 Hz. Signal conversion was performed using custom-written MatLab (version 2019a; The MathWorks Inc, Natick, Massachusetts, USA) scripts and the output was converted into European Data Format for offline analysis. For each 24 h recording, EEG power spectra were calculated via Fast Fourier Transform (FFT) for 4 s epochs, at a 0.25 Hz resolution (SleepSign Kissei Comtec, Nagano, Japan).

### Detection of hypothermia bouts

In this study, T_surface_ was recorded using thermal imaging cameras, and used for detecting the occurrence of hypothermia bouts, defined by transient decreases of T_surface_ (see below). We noted that the absolute values of T_surface_ obtained in this study (**Fig. 1**) were consistently lower than previously reported temperature values recorded in small rodents from the brain or intraperitoneally placed thermistors. Therefore, the values of T_surface_ reported here cannot be interpreted as a proxy of core body or brain temperature, especially that T_surface_ recordings are likely more prone to artifacts, arising from movement and changes in the visibility of the “hotspot” to the cameras depending on the animal’s position (**Fig. 1**). Transient decreases in T_surface_ occurred in all 6 animals, and were initially superficial, e.g. the drop in T_surface_ was 1–2°C below the median temperature during baseline. As fasting progressed, the bouts of hypothermia became progressively deeper and longer, and unequivocal torpor bouts occurred in all animals on Day 5 of fasting (representative example: **Suppl. Fig. 1a**). Hypothermic bouts were detected using custom-made Matlab scripts based on T_surface_ data averaged in 1 min bins and smoothed with a 20 min moving average, which removed artefacts, occurring as a result of movement. During days of fasting, hypothermia bouts were defined as time periods during which T_surface_ was more than 3 SD below the median temperature value, and which ended when T_surface_ reached at least the level of 1 SD below the mean T_surface_ recorded on baseline days. Upon inspection, it was revealed that such periods sometimes consisted of “sub-bouts” that were demarcated by noticeable increases in T_surface_, while remaining well below euthermic levels. These “sub-bouts” were not considered as interruptions of hypothermia bouts. Substantial variability was observed across hypothermia bouts with respect to minimal T_surface_ achieved. To allow comparison between baseline and fasting days (including the time intervals after the animals were fed on the days with food restriction), for subsequent analyses we identified all time periods when T_surface_ declined by at least 0.5°C relative to median T_surface_ calculated over baseline days. To investigate the relationship between T_surface_, vigilance states and the EEG spectra, we further identified epochs of “deep hypothermia” on the days of food restriction, where T_surface_ was decreased by more than 4 °C relative to the median T_surface_ calculated at baseline.

### Scoring of vigilance states

Scoring of vigilance states was performed offline by visual inspection of consecutive 4 s epochs (SleepSign, Kissei Comtec, Nagano, Japan). Frontal and occipital EEG derivations and EMG were displayed simultaneously to facilitate scoring. Vigilance states were classified as wake (high frequency, low-amplitude irregular EEG pattern dominated by theta-activity, 6–9 Hz), non-rapid eye movement sleep (NREM; EEG dominated by high amplitude, low frequency waves), or rapid eye movement sleep (REM; EEG is dominated by theta-activity, most prominent in the occipital derivation, with a low level of EMG activity). Epochs where EEG signals were contaminated by artefacts due to movement were excluded from spectral analyses (7.1±3.2% of total recording time). The onset of individual NREM sleep episodes was defined by the first occurrence of slow waves in at least one EEG channel, along with the absence of EMG activity. Vigilance states annotation was performed across all days, including the time periods when T_surface_ was low (see Results). As EEG amplitude decreased in association with a drop in T_surface_, it was not used as a key criterion for vigilance state annotation and the scoring was based on frequency content and the overall pattern of EEG activity, in addition to the presence or absence of EMG tone. To incorporate the possibility that vigilance states, including NREM sleep, in the euthermic condition are inherently different from those during hypothermia, we performed separate analyses of their amounts and corresponding EEG spectra for euthermic and hypothermic epochs.

### Statistics

Statistical analyses were performed using Microsoft Excel 365 (Microsoft, USA)^33^ or Matlab (The MathWorks Inc, Natick, Massachusetts, USA). Since EEG spectral power values are not normally distributed, data were log-transformed prior to statistical comparison.^34^ Data are presented as mean values with standard error of the mean (SEM). To assess the effect of fasting across days, one-way repeated measures ANOVA was used. Pair-wise comparisons were calculated based on parametric (paired Student’s T) tests.

## Results

### Body weight and temperature

Body weight and T_surface_ were recorded throughout the experiment. As restricted feeding progressed, a decrease in mean body weight was observed. Over 5 days of restricted feeding, mean body weight fell from 28.3 ± 1.17 g during baseline to 24.1 ± 0.94 g on Day 5 (p=0.00002; F(2, 10) = 233.99, p = 4.0081e-09, repeated measures ANOVA; **Fig. 1c**). Over the same time period, daily T_surface_ values also decreases: maximum T_surface_ decreased slightly from 32.6 ± 0.24 °C to 31.4 ± 0.075 °C (p=0.0049; F(5, 25) = 12.5, p = 3.9393e-06), mean T_surface_ dropped from 31.6 ± 0.27 °C to 27.2 ± 0.47 °C (p=0.0014; F(5, 25) = 17.7, p = 1.6608e-07), whilst minimum T_surface_ dropped substantially from 30.1 ± 0.36 °C to 21.5 ± 0.61 °C (p=0.0002; F(5, 25) = 25.9, p = 3.8435e-09) (**Fig. 1d**). Overall there was a negative correlation between body weight and mean daily T_surface_ (**Fig. 1e**).

### Characteristics of hypothermic bouts

From T_surface_ data, hypothermic bouts, e.g. any decreases of peripheral temperature by > 0.5°C relative to median baseline value, were detected on all days (See Methods). As fasting progressed, the bouts of hypothermia become progressively longer and deeper (**Fig. 2a**). Calculating the incidence of hypothermia bouts across consecutive days revealed that the number of bouts per 24h did not change significantly and averaged 3.8 ± 0.91 on Day -1 and 4.7 ± 0.61 on Day 5 (F(5, 25) = 1.2, n.s.; **Fig. 2b**). However, a marked increase in mean duration of hypothermia bouts was evident, starting from 121 ± 12 min on Day -1 and reaching 213 ± 37 min on Day 5 (p=0.0240; F(5, 25) = 5.1, p=0.002; **Fig. 2c**). The minimum temperature values attained during hypothermia bouts also decreased progressively from 30.2 ± 0.31 °C on Day -1 to 25.0 ± 0.52 °C (p=0.0008; F(5, 25) = 14.9, p=8.009e-07) on Day 5 (**Fig. 2d**). This was reflected in a shift towards a more frequent occurrence of hypothermia bouts with lower values of T_surface_ during days of fasting as compared with baseline days when food *ad libitum* was provided (p=0.006; **Fig. 2e**). Next, we calculated an index of hypothermia “intensity” ^25^, by integrating all temperature values (expressed as a decrease relative to median baseline T_surface_) during each hypothermia bout. This analyses revealed a progressive increase of hypothermia index across days (F(5, 25) = 5.7, p=0.012; **Fig. 2f**), and a greater incidence of hypothermia bouts, characterised by their higher intensity, during fasting, as compared to food *ad lib* condition (p=0.021; **Fig. 2g**).

**Figure 2.**
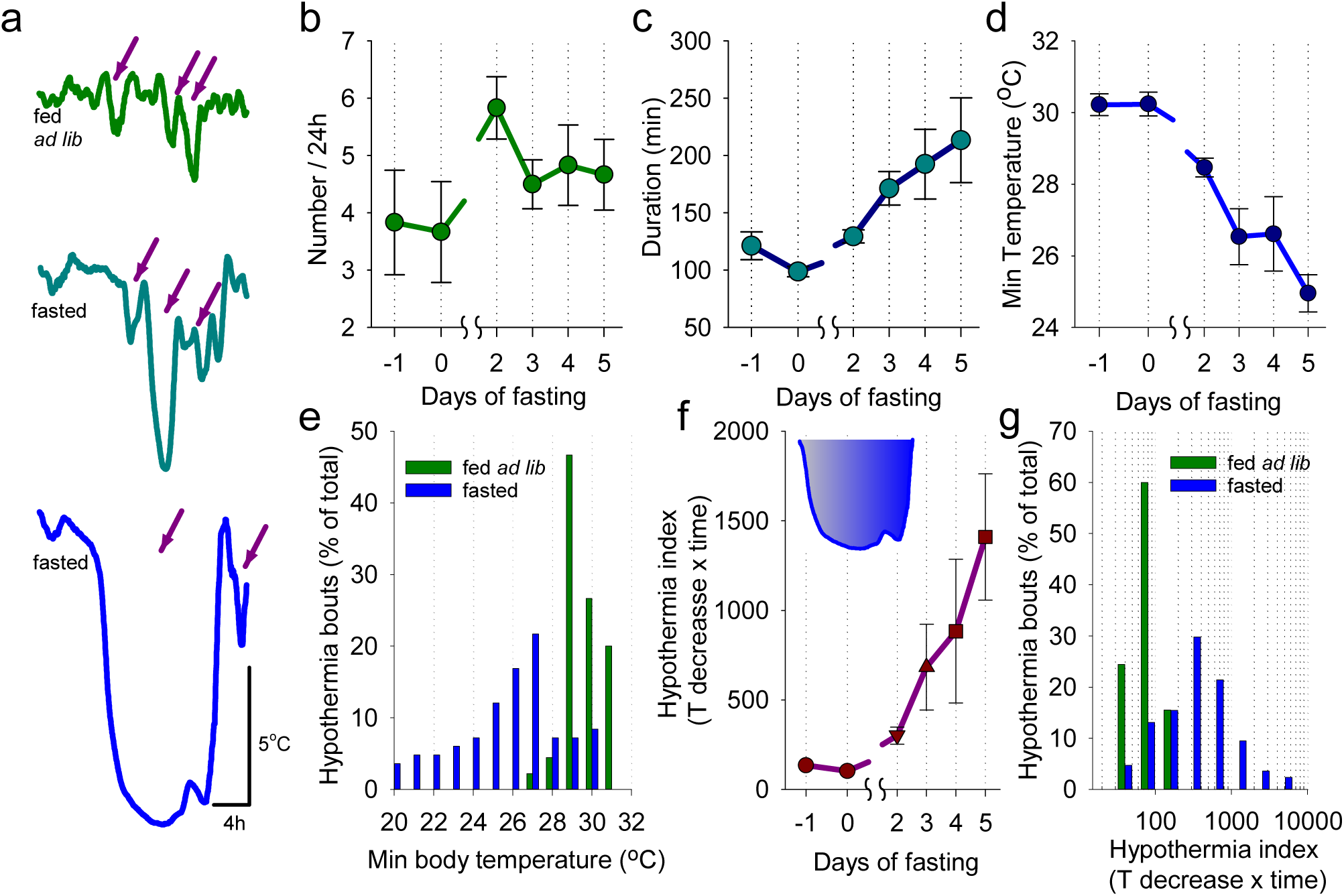
The characteristics of hypothermia bouts. **a**, Representative examples of hypothermia bouts during baseline (fed *ad lib*) and on the 3^rd^ and 5^th^ days of food restriction. Arrows depict bouts of hypothermia used for subsequent analyses. **b**,**c**,**d**, The time course of the number and duration of hypothermia bouts and the minimal surface body temperature attained during hypothermia bouts. Note that as restricted feeding progressed, hypothermia bouts increased in duration, became more frequent and deep. N=6, mean values, SEM. **e**, Distribution of hypothermia bouts during baseline and fasted days. **f**, The time course of hypothermia index (a variable taking into account both depth and duration of hypothermia; represented by inset showing shaded area between curve and straight line) across the experiment. **g**, Distribution of hypothermia bouts as a function of hypothermia index. Note that as restricted feeding progressed, the hypothermia index increased.

### The relationship between hypothermia and vigilance states during fasting

Next, we evaluated the temporal pattern of hypothermia bouts occurrence. During baseline, superficial bouts of hypothermia occurred generally uniformly across 24h **(Fig. 3a,b)**. However, as fasting progressed, deep bouts of hypothermia occurred most prominently towards the middle of the dark period and recurred prior to the feeding interval **(Fig. 3a,b)**. The visual inspection of EEG spectra revealed that EEG power was generally depressed across all frequencies during hypothermia bouts, especially when T_surface_ was significantly reduced **(Fig. 3c)**. However, the typical EEG and EMG signatures of wakefulness, NREM sleep and REM sleep were apparent, which allowed performing vigilance state annotation throughout the recording period **(Fig. 3d)**. We observed that as fasting progressed, the amount of wakefulness as expressed as a percentage of 24 h increased initially from 50.2 ± 2.4% to 59.7 ± 2.3% (p=0.0017) on Day 3, but then decreased on Day 5 to 51.3 ± 4.7% (p=0.1800, F(6, 30) = 2.5, p=0.042; **Fig. 4a, top**). At the same time, the daily amount of NREM sleep (including epochs during both euthermia and hypothermia) showed first a suppression but then returned to values similar to baseline (fed ad lib: 41.6 ± 1.8%, Day 5: 47.1 ± 4.7%, p=0.29; F(6, 30) = 4.9, p=0.001; **Fig. 4a, middle**). However, the amount of REM sleep decreased markedly from 8.1 ± 0.5% at baseline to 1.6 ± 0.2% (p=0.0001) on Day 5 (F(6, 30) = 35.6, 2.4163e-12; **Fig. 4a, bottom**). These changes were evident from individual hypnograms (three days from a representative individual mouse shown on **Fig. 4b**). Next, we subdivided epochs scored as NREM sleep into those occurring at euthermia and those that occurred during hypothermia. This analysis revealed that the amount of NREM sleep during euthermia periods decreased significantly from 28.0 ± 3.8% on Day -1 to 7.9 ± 1.5% on Day 5 (p=0.0048, F(5, 25) = 14.4, p=1.0805e-06), while the amount of hypothermic NREM sleep showed an opposite trend from 13.6 ± 2.4% on Day -1 to 39.2 ± 5.0% on Day 5 (p=0.0001, F(5, 25) = 15.6, p=5.5161e-07, **Fig. 4c**). These findings show that as fasting progresses, NREM episodes become more prevalent during hypothermic bouts. Focusing specifically on the proportion of each vigilance state within hypothermic bouts on Day 5 and during matching time periods on Day -1, revealed lower amounts of wake (31.2 ± 3.8% versus 45.6 ± 2.1%; p=0.012) and REM sleep (0.96 ± 0.36% versus 8.2 ± 0.5%; p=0.00004), while the amount of NREM sleep was increased (61.8 ± 3.4% versus 42.5 ± 1.5%; p=0.002). The decrease in T_surface_ was strongly associated with the amount of REM sleep, which was proportionally decreased, while NREM sleep increased as a function of hypothermia deepening (**Fig. 4d**). The T_surface_ and the amount of wakefulness were only weakly related.

**Figure 3.**
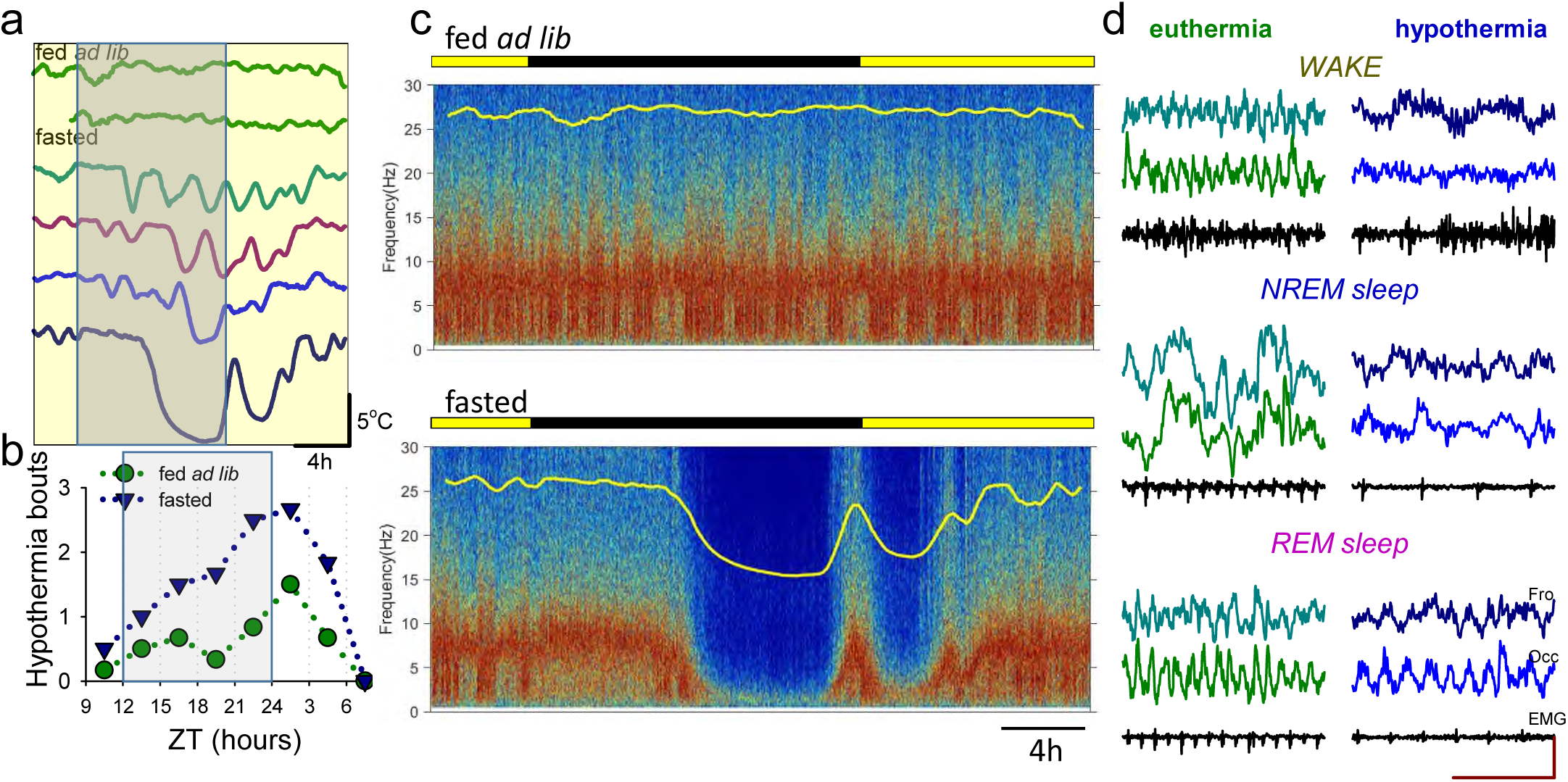
The timing of torpor bouts and corresponding changes in the EEG. **a**, Representative example of body temperature in one individual animal showing that deep bouts of hypothermia during fasting are typically clustered toward the middle of the dark period. **b**, The time course of hypothermia bout occurrence across 24h during baseline (fed ad lib) and during fasting. **c**, Representative EEG spectrograms during baseline day and during the last day of fasting in one individual mouse. Note a substantial reduction in EEG power during hypothermia. **c**, Representative EEG and EMG traces taken from wakefulness, NREM sleep and REM sleep in euthermic condition (surface body temperature > 30°C) and during hypothermia (< 24°C).

**Figure 4.**
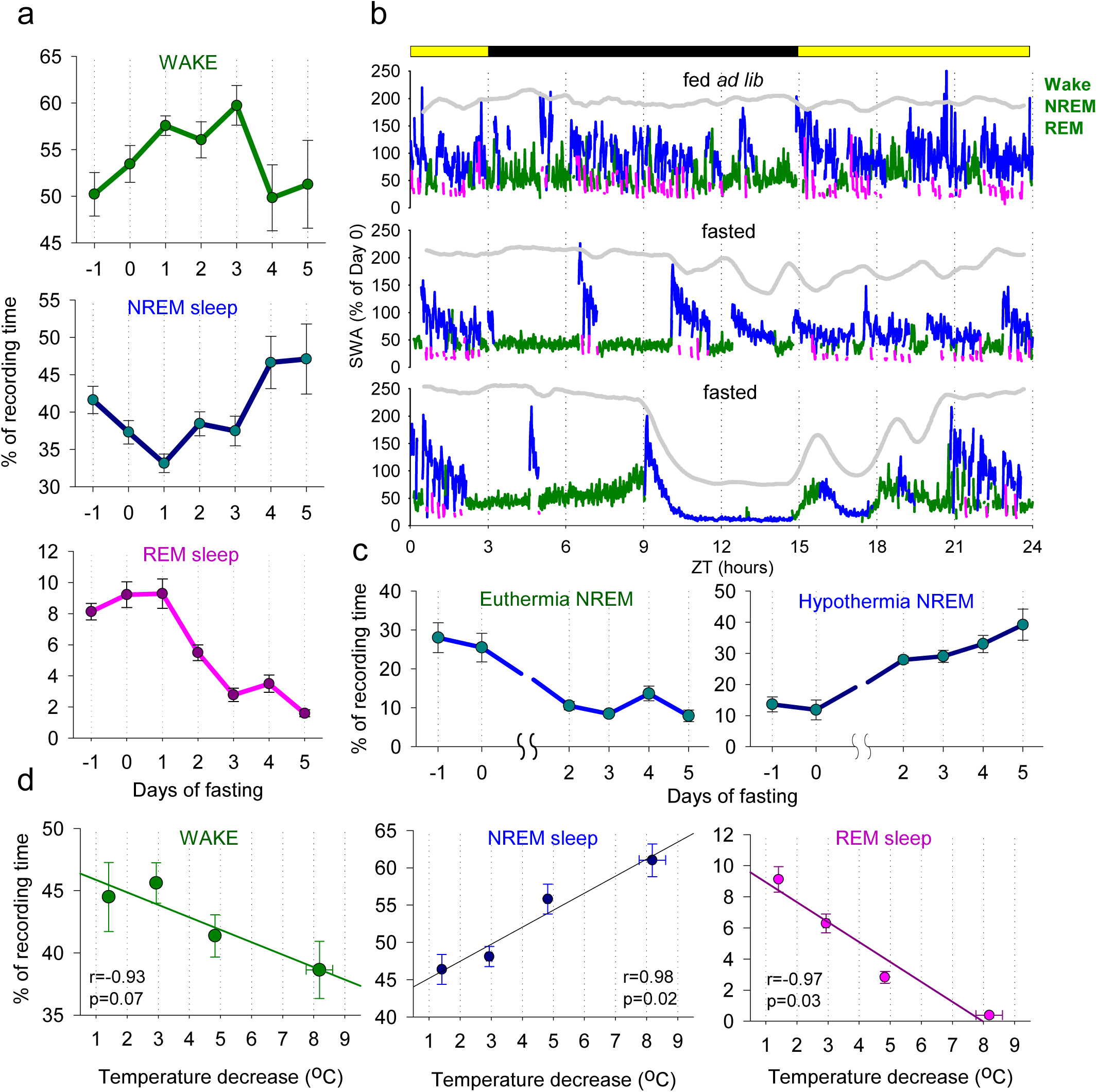
The effects of fasting and hypothermia on vigilance states. **a**, The time course of daily amount of EEG/EMG defined wakefulness, NREM sleep and REM sleep. N=6, mean values, SEM. **b**, The time course of EEG slow-wave activity (0.5-4Hz, SWA) during the 24h period shown for baseline day and two days of food restriction (Day 3 and Day 5) in one individual mouse. Mean SWA is plotted in 1-min epochs and is color-coded according to the vigilance state (waking: green, NREM sleep: blue, REM sleep: pink). The curve at the top is corresponding to surface body temperature. Note the drop in SWA when body temperature is low. **c**, The proportion of NREM sleep during euthermia and during hypothermia bouts. Note a progressive increase in the amount of NREM sleep during fasting. N=6, mean values, SEM. **d**, The relationship between the amount of waking, NREM and REM sleep and peripheral body temperature. Note that when the temperature declines by more than approximately 5°C, REM sleep virtually disappears.

### EEG spectral analysis during wake and sleep: effects of hypothermia

Next, we investigated the effects of fasting and hypothermia on EEG spectral power. To this end, we addressed whether the decrease in spectral power we observed (**Fig. 3c**) was state specific and whether it was primarily associated with T_surface_ or resulted from changes in sleep intensity associated with fasting. We calculated EEG power spectra separately for epochs, scored as waking, NREM sleep and REM sleep during baseline when the animals were fed *ad libitum*, during epochs of deep hypothermia when the animals were fasted, and also during those epochs on fasting days when T_surface_ was similar to baseline. We observed that EEG power generally showed a decrease during hypothermia on days when the animals were food restricted, but it was virtually identical between euthermia epochs on fasted days and during baseline (**Fig. 5a**). The reduction in EEG power during hypothermia was especially pronounced during NREM sleep, which can be expected based upon the predominant occurrence of this state during hypothermia bouts (**Fig. 4c**). The decrease in EEG power during waking was also observed during hypothermia as compared to both baseline and euthermia in the frontal derivation and compared to euthermia only in the occipital EEG (**Fig. 5a**). The decrease in EEG power during REM sleep was somewhat more pronounced than during waking, but caution is warranted with interpreting this result as the total amount of REM sleep was drastically decreased when T_surface_ was low. However, during those epochs of REM sleep that occurred during hypothermia, a marked left-ward shift of the theta peak was present, consistent with the observation made previously in Djungarian hamsters.^35^ To further address whether the changes in the EEG observed were related to temperature changes rather than fasting, we clustered all waking, NREM and REM-scored epochs as a function of progressively decreasing T_surface_, and calculated corresponding total spectral EEG power in the frequency range between 0.5-30Hz (**Fig. 5b**). As expected, we observed generally higher values of total EEG power during NREM sleep at euthermia, but in all three vigilance states EEG power decreased markedly as a function of T_surface_ decrease (**Fig. 5b**).

**Figure 5.**
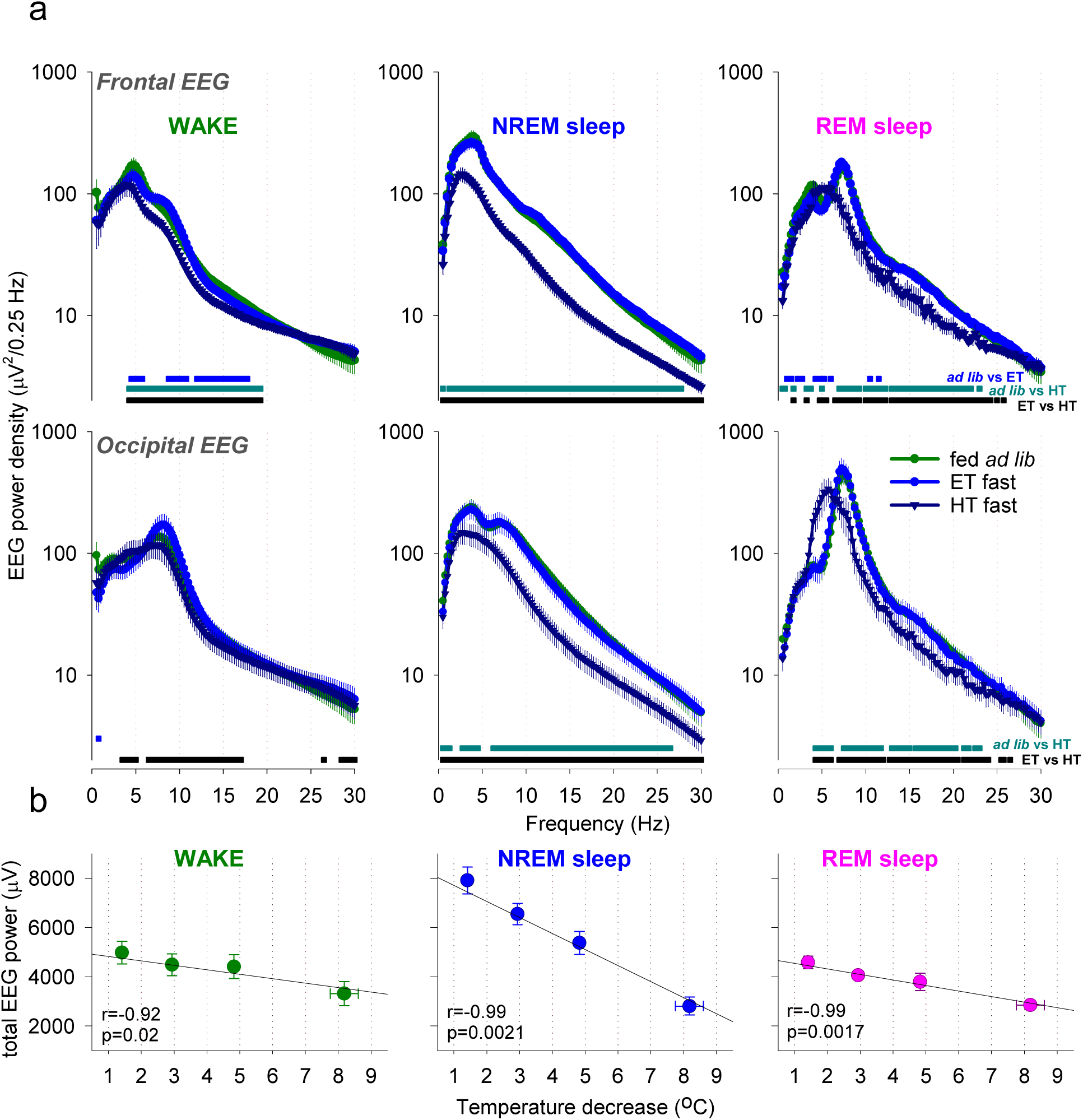
The effects of fasting and hypothermia on EEG power spectra. **a**, EEG power spectra during waking, NREM sleep and REM sleep during baseline (fed *ad lib*) and shown separately for euthermic (ET) and hypothermic (HT) episodes during fasting. Note that EEG power generally declines during hypothermia on fasted days, but is virtually indistinguishable from spectra of the EEG recorded during the same days at euthermia. EEG power spectra during REM sleep highlight a marked slowing of theta peak frequency. N=6, mean values, SEM. **b**, The relationship between total EEG power during waking, NREM and REM sleep and peripheral body temperature. Note a strong negative relationship between peripheral body temperature and EEG power in all vigilance states.

### Hypothermia bouts are initiated from deep NREM sleep

The visual inspection of individual hypnograms suggested that bouts of hypothermia do not start from wakefulness or REM sleep, but rather commence during NREM sleep with high SWA (**Fig. 4b,6a**). Notably, during the initial NREM sleep at the beginning of hypothermia bouts, the EEG signals were indistinguishable between those that progressed into deep hypothermia or normal NREM sleep associated with a minor decrease in T_surface_ only (**Fig. 6b**). To systematically evaluate this observation, we identified all hypothermia bouts lasting at least 2 h and calculated the corresponding amount of sleep. For these analyses, we grouped all hypothermic bouts into those where T_surface_ showed only a minor decrease of not more than 2 °C, and those that progressed into deep hypothermic bouts (decrease more than 4 °C from baseline; **Fig. 6c**). We observed that, in both cases, the onset of hypothermia was associated with a marked increase in the proportion of NREM sleep, which was especially pronounced at the onset of deep hypothermic bouts (**Fig. 6d**). Furthermore, as expected, a greater amount of NREM-like state was seen as T_surface_ decreased further. At the same time, EEG SWA started with high values in both cases, and showed a similar decreasing trend during the following 60 min period (**Fig. 6e**). Thus, these data suggest that the occurrence of bouts of hypothermia is closely linked to the occurrence of deep NREM sleep characterised by high SWA.

**Figure 6.**
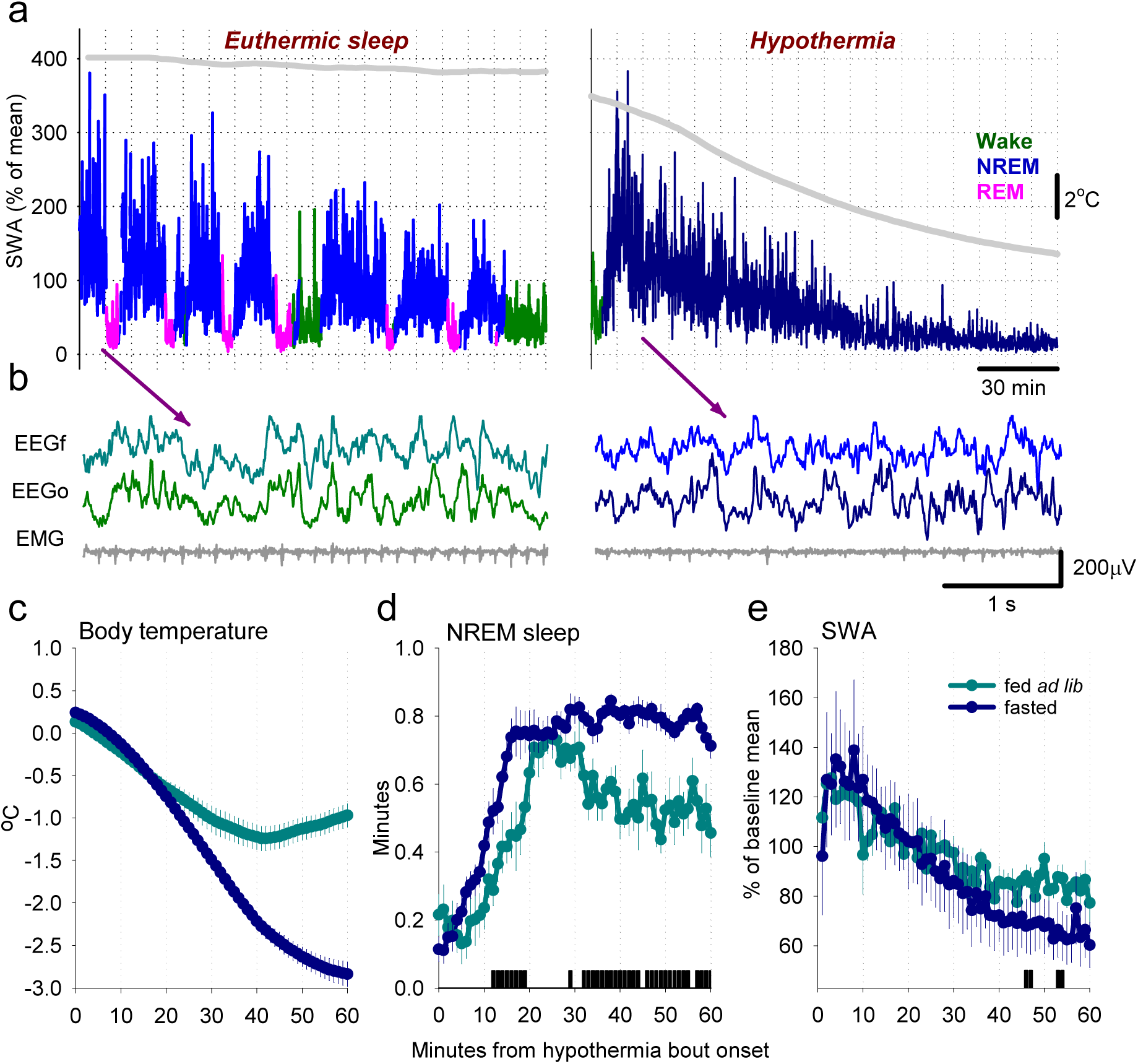
The relationship between surface body temperature, sleep and SWA at the onset of hypothermia episodes. **a**, The time course of EEG slow-wave activity (0.5-4Hz, SWA) during a typical period of sleep at euthermia and the dynamics of SWA during the entrance into a deep bout of hypothermia. SWA is plotted in 4-s epochs and is color-coded according to the vigilance state (waking: green, NREM sleep: blue/dark blue, REM sleep: pink). The curve at the top is corresponding to surface body temperature. Note the drop in SWA in both cases, but it is especially pronounced as the temperature decreases. **b**, Representative EEG traces of NREM sleep at the beginning of sleep periods when the surface body temperature remains high or subsequently declines. Note that during this time EEG activity is virtually indistinguishable, suggesting that even deepest hypothermia bouts start from a NREM sleep state. **c,d**, The time course of peripheral body temperature and the proportion of NREM sleep starting from the onset of a hypothermia bout in animals fed *ad lib* and fasted animals. N=6, mean values, SEM. Note the rapid increase in the amount of NREM sleep at the beginning of a hypothermia bout in fasted animals and a greater proportion of NREM sleep later during the hypothermia bout. The bar on the bottom denote significant differences (p<0.05, paired t-test). **e**, The time course of EEG SWA during NREM sleep from the onset of hypothermia bout in fed and fasted animals. Note that the values of SWA at high and show a progressive decrease in both cases. N=6, mean values, SEM.

### Bursts of EMG activity during hypothermia

The EMG activity dropped rapidly at the very beginning of the hypothermia bouts, at a markedly higher rate than the decrease of T_surface_ (**Fig. 7a**), as could be expected from the predominant occurrence of a NREM sleep like state during this time (**Fig. 6d**), although residual EMG tone was still initially present. Plotting individual spectrograms alongside with EMG activity on the day 5 of fasting indicated an unexpectedly high level of EMG activity during bouts of deep hypothermia, especially later in their progression (**Fig. 7b**). A close inspection revealed that EMG activity is not tonic and continuous but occurs in a form of regularly occurring discharges, sometimes happening with a striking periodicity (**Fig. 7c,d**). EMG bursts typically started with an occurrence of a high amplitude EEG potential, and lasted between two and three 4 s epochs only, during which the EEG was activated (**Fig. 7e**). To analyse the occurrence of EMG bursts in more detail, we selected one bout of hypothermia in each animal, in all cases occurring during the last day of restricted feeding. To detect EMG bursts, we used an individually determined threshold which was applied to consecutive values of EMG variance calculated based on 4 s epochs, and the onset and the end of all events lasting less or equal than 20 s were calculated (the average EMG profile centred on the starting epoch of EMG bursts shown on **Fig. 7f**). The majority of inter-burst intervals were around 2 min (on average 2.3±0.3 min), although more frequent occurrence of EMG bursts, or several minutes long periods without EMG discharges, were not uncommon (**Fig. 7g**). Finally, we calculated the incidence of EMG bursts during the period of hypothermia-associated immobility, which revealed a progressive 3 to 4-fold increase in the occurrence of EMG bursts, which occurred in parallel with the decrease in T_surface_ (F(19, 95) = 4.1, p= 2.4336e-06, **Fig. 7h**). Thus, hypothermia bouts do not correspond to a behavioural state with a total depression of EMG tone, but are characterised by a regular occurrence of motor discharges.

**Figure 7.**
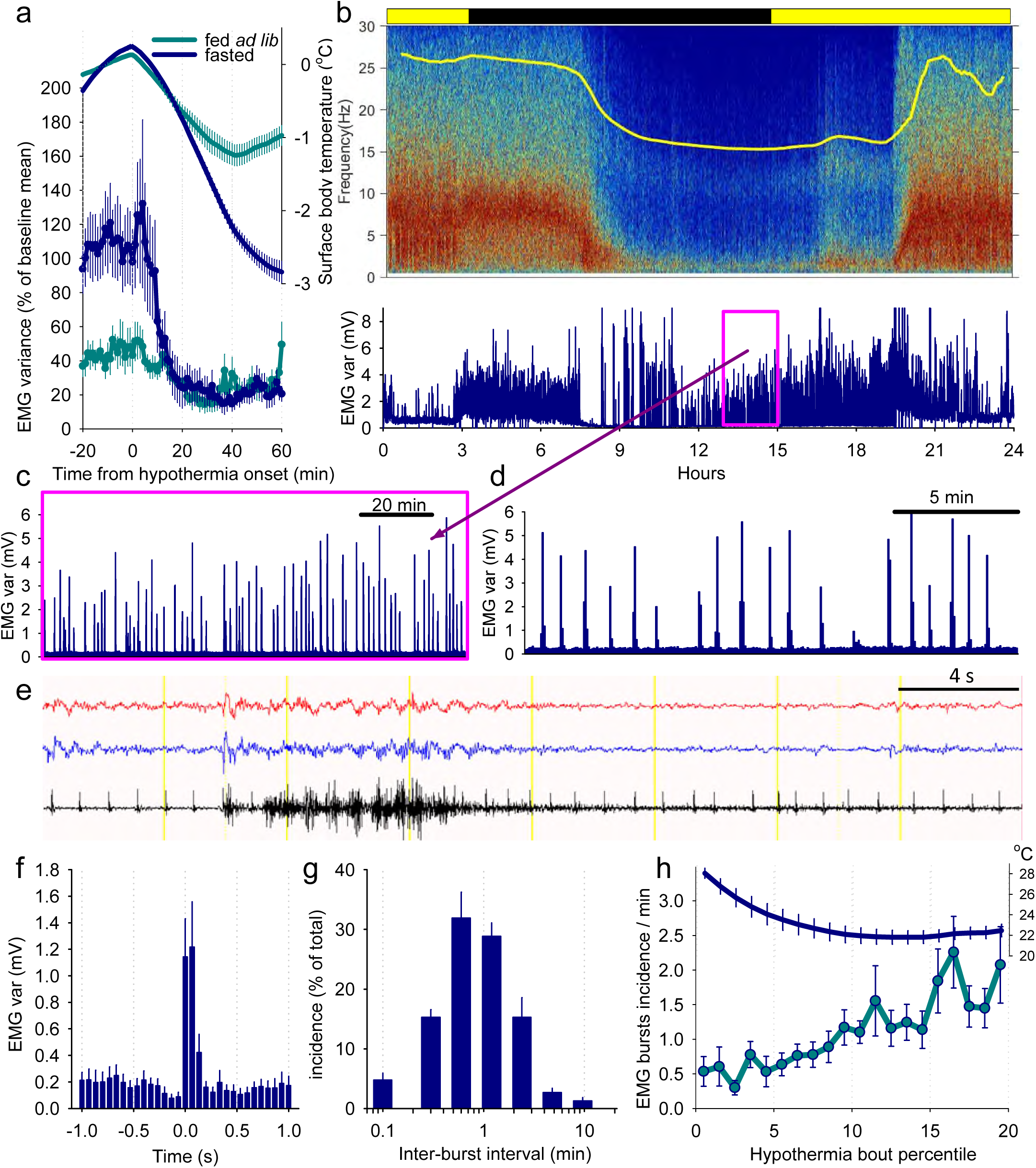
Periodic bursts of muscle activity during hypothermia bouts. **a**, The time course of EMG variance and surface body temperature aligned to the onset of hypothermia bouts in fed *ad lib* and fasted mice. Mean values, N=6, SEM. **b**, EEG spectrogram of one representative animal before, during and after a prolonged bout of hypothermia. The panel below depicts corresponding EMG variance. **c**, The 2-h interval outlined in a box on panel **b** is shown at a greater temporal resolution. **d**, The same focused on a shorter 20-min window. Note the occurrence of regular bursts of EMG activity. **e**, Representative EEG and EMG traces recorded during hypothermia highlighting an occurrence of a single EMG burst. **f**, Average profile of EMG variance centered on the onset of individual EMG bursts. **g**, Distribution of inter-burst intervals. **h**, The time course of EMG burst incidence and corresponding body temperature between the onset and offset of hypothermia related immobility. N=6, mean values, SEM. Note that EMG bursts become more frequent as temperature declines.

## Discussion

This is the first study to perform a detailed investigation of EEG/EMG defined states of vigilance during hypothermia and torpor induced following restricted feeding in mice. The observation that fasting in the laboratory mouse induces progressively deeper bouts of hypothermia confirms several previous studies.^26,27,32,36^ Although EEG has been recorded during hibernation in the Arctic ground squirrel, in gray mouse lemurs^37^ and during daily torpor in the Djungarian hamster, ^38^ how brain activity changes during fasting-induced torpor in the laboratory mouse has been under-investigated.

Torpor is primarily defined as a state characterised by a reduced metabolic rate ^13,39^. By contrast, sleep and wake states are typically classified based upon characteristic patterns of brain activity.^40^ Here we observed that as peripheral body temperature decreased and torpor progressed, EEG power dropped substantially, yet distinctive EEG and EMG signatures of euthermic wakefulness and sleep were not completely lost. The extent to which this hypothermic sleep-like state is comparable to euthermic sleep, both neurophysiologically and functionally, remains to be resolved. However, this study provides novel insights which help addressing the existing controversy regarding the relationship between torpor and sleep.

We observed that as the restricted feeding paradigm progressed, the prolongation and deepening of hypothermic periods were associated with an increase in the amount of epochs scored as NREM sleep. Whether this hypothermia-associated NREM-like state alleviates homeostatic sleep pressure and, if so, to what degree remains an intriguing open question. It is possible that sleep debt accumulates during fasting-induced torpor, similar to daily torpor in Djungarian hamsters.^41^ However, this remains to be addressed in future studies.

We also observed a marked decrease in REM sleep, both with successive days of fasting and with successively lower body temperatures. It appears that REM sleep is essentially abolished below a T_surface_ of 25 — 26 °C, consistent with previous studies demonstrating the temperature dependence of REM sleep.^42,43^. As expected, the amount of waking initially increases (up to Day 3), possibly reflecting the increase in arousal and food-seeking behaviours, ^6,44^ before decreasing when the energy saving strategy of torpor is fully established by Day 5 of food restriction.

A novel insight from our study was the observation that entrance into a hypothermic bout invariably coincides with a transition from the awake state to a state indistinguishable from NREM sleep. This transition preceded and was much faster than the gradual decline in T_surface_. Adenosine signalling in the hypothalamic preoptic area (POA) has been implicated in the central control of both sleep regulation and thermoregulation. Due to this similarity, it would not be a surprise if torpor were entered via a state resembling NREM sleep.^45–48^ A possible interpretation is that due to the close proximity of centres involved in regulating sleep and body temperature, adenosine signalling that results from a pro-hypometabolic signal can “spill over” from the temperature control centre to the sleep control centre. ^49–51^ An intriguing observation is that whilst going into torpor, animals have to overcome the particularly strong wake drive associated with hunger in order to implement a longer term strategy of energy saving.

As T_surface_ decreases upon torpor onset, the levels of SWA progressively decreased as well. These changes were gradual and therefore it is difficult to precisely define a time point at which the EEG no longer resembles typical euthermic NREM sleep. It remains unclear whether the decrease in EEG power during hypometabolism is merely an epiphenomenon directly related to a reduction in brain temperature, which would be expected to slow down all relevant biochemical and cellular processes and synaptic neurotransmission ^21^. Consistent with this view is our finding of a positive correlation between T_surface_ and EEG spectral power. It is also possible that other changes contribute to the reduction of SWA. For example, structural changes in synapses, activation of presynaptic and postsynaptic adenosine receptors resulting in reduced neuronal excitability, closure of temperature-sensitive TRPV3 and TRPV4 channels, and downstream effects of cold-inducible transcription factors such as RBM3 could all play a role ^22,23,52–55^. Facilitated by higher ambient temperatures in their natural habitats, some animals (e.g. the Madagascan lemur) naturally undergo hypometabolism at their euthermic body temperature. Whether under these circumstances there are EEG changes similar to those we have described, or whether these changes are a consequence of hypothermia remains to be determined.

Unlike in extrinsically-induced hypothermia, shivering thermogenesis in mice entering torpor is suppressed.^56–58^ As a torpor bout progressed, T_surface_ eventually reached a stable level at approximately 1-2 °C above ambient temperature. During this “maintenance” phase of torpor, we observed periodic bursts of EMG activity, typically lasting 8-12 seconds. The EMG bursts were especially regular during the lowest levels of hypothermia. One possibility is that they represent brief episodes of shivering thermogenesis, involuntary somatic motor response thought to be mediated by spinothalamic afferents and the median preoptic nucleus.^56–58^ This may allow defence of core body temperature above that of ambient temperature.^21,26^ Intriguingly, the occurrence of EMG bursts also resembles the regular motility patterns typical of sleep during early ontogeny. ^59–61^ Although the function of motor discharges during development remains unclear, it has been proposed that they may also reflect a default mode of activity typical of the immature nervous system. Since it can be expected that, during deep hypothermia, most brain functions are deeply depressed, we surmise that this may have unmasked an emergence of primitive, stereotyped patterns of activity driven by the central pattern generator. The rewarming phase of torpor was characterised by a marked increase in the frequency and intensity of these bursts, and the rate of rewarming was observed to be faster than that of entry. This is likely due to the latter being a passive process (i.e., conduction and radiation of heat from the body to the environment), and the former being an active process involving increased shivering and non-shivering thermogenesis.^56–58^

In summary, our data suggest that on the basis of electrophysiogical recordings, torpor and sleep are not necessarily mutually exclusive states. Rather, we suggest that hypometabolism and associated hypothermia characterized by suppressed EEG activity and periodic motor discharges, shares electrophysiological characteristics with sleep. Whether and how, sleep associated with fasting-induced torpor is homeostatically regulated remains to be determined.

## Acknowledgments

Supported by: Guarantors of Brain Entry Clinical Research Fellowship, MRC and Stroke Association Jointly-Funded Clinical Research Training Fellowship (MR/S001948), Wellcome Trust Strategic Award (098461/Z/12/Z), Wellcome Trust Senior Investigator Award (106174/Z/14/Z), John Fell OUP Research Fund Grant (131/032). We thank M. Guillaumin, L. Krone, L. Milinski, L. McKillop, C. Blanco-Duque and M. Kahn for help with experiments.

---

**Suppl. Figure 1.**
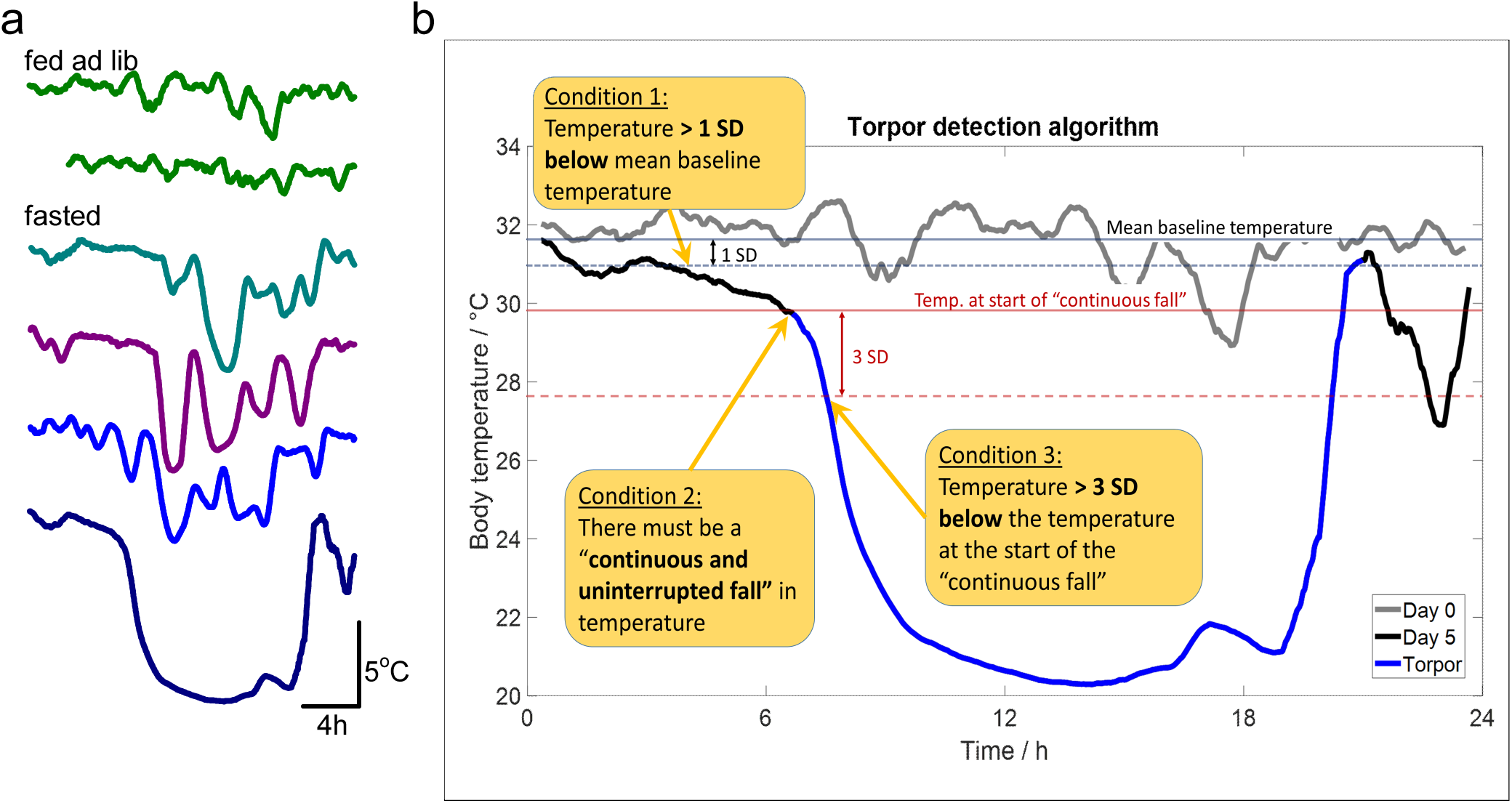
Detection of hypothermia bouts during baseline and food restriction. **a**, Representative example of body temperature in one individual animal showing that bouts of hypothermia may occur during baseline in fed ad lib animals, but become more frequent, longer and deeper as fasting progresses. **b**, Schematic showing how hypothermia bouts were detected using an algorithm, based on a heuristic method of identifying a set of thresholds for determining the onset and end of individual hypothermia bouts. In all cases the detections were checked visually and corrected where it was deemed necessary.

